# Evaluation of habitat protection under the European Natura 2000 conservation network – the example for Germany

**DOI:** 10.1101/359125

**Authors:** Martin Friedrichs, Virgilio Hermoso, Vanessa Bremerich, Simone D. Langhans

**Author notes:** Corresponding author: Martin Friedrichs.

## Abstract

The world’s largest network of protected areas - Natura 2000 (N2000) - has been implemented to protect Europe’s biodiversity. N2000 is built upon two cornerstones, the Birds Directive, which lists 230 bird species and the Habitats Directive, which lists next to a variety of species 230 habitat types, to be protected. There is evidence of the positive impact of the Directives on the EU’s biodiversity, although the overall improvement reported for species in favourable condition in the last assessment was low. However, most of the assessments are species focused, while habitats have received very little attention. Here we developed a generic workflow, which we exemplified for Germany, to assess the status of habitat coverage within the N2000 network, combining information from publicly available data sources. Applying the workflow allows identifying gaps in habitat protection, followed by the prioritization of potential areas of high protection value, using the conservation planning software Marxan. We found that, in Germany, N2000 covers all target habitats. However common habitats were proportionally underrepresented relative to rare ones, which contrasts with comparable studies for species. Moreover, the German case study suggests that especially highly protected areas (i.e. covered by more than 90% with N2000 sites) build an excellent basis towards a cost-effective and efficient conservation network. Our workflow provides a generic approach to deal with the popular problem of missing habitat distribution data outside of N2000 sites, information which is however crucial for managers to plan conservation action appropriately across Europe. To avoid a biased representation of habitat types within N2000, our results underpin the importance of defining qualitative and quantitative conservation targets which will allow assessing the trajectory of habitat protection in Europe as well as adjusting the network accordingly - a future necessity in the light of climate change.

## 1. Introduction

In times of global biodiversity loss (Cardinale et al., 2012), large nature conservation networks will become increasingly important for mankind, as human wellbeing is closely linked to biodiversity (Diaz et al., 2006). The most important contribution to biodiversity conservation in Europe is Natura 2000 (N2000), the world’s largest nature conservation network. N2000 is built upon two cornerstones: The Birds Directive (BD), introduced in 1979 (Council Directive 79/409/EWG) was established to protect bird species, their resting areas, and migration pathways through the implementation of special protection areas (SPAs). Since 1994, these SPAs are included in the N2000 network. The second cornerstone of N2000 is the Habitats Directive (HD; Council Directive 92/43/EEC), introduced in 1992, which is based on the designation of special conservation areas (SCAs) that should ensure the protection of a broader range of species. In the Annexes II, IV, and V, more than 1000 animals and plants are listed which need protection across the EU. Additionally, Annex I lists more than 230 rare and characteristic habitat types which must be considered.

The HD can be seen as the backbone of habitat conservation efforts in Europe and does not have a counterpart in the rest of the world (Hodge et al., 2015). The explicit effort to protect habitats is a reaction to the increasing awareness that, besides species, also habitats are under great threat and essential for successful nature conservation (e.g. European Red List of Habitats; IUCN, 2017). It is well known that an increased habitat diversity is beneficial for species diversity (Hortal et al., 2009). Additionally, habitats are commonly used as biodiversity surrogates in conservation assessments (Bunce et al., 2013). Therefore, an evaluation of habitat coverage under N2000 is a good reflection of how well N2000 is covering biodiversity overall.

Despite a worldwide commitment, human pressures and inadequate conservation efforts limited the success of reducing the rate of biodiversity loss by 2010 (Butchart et al., 2010) and new targets were set to 2020 (CBD, 2017). These targets mandate that 17% of terrestrial area and 10% of coastal and marine areas should be covered by nature conservation networks (Target 11). Furthermore, by 2020 100% more habitat assessments should show an improved conservation status compared to current assessments (Target 1, EU Biodiversity Strategy to 2020). Considering that N2000 currently protects 18% of terrestrial and 6% of Europe’s marine area (European Commission, 2016a) the BD and HD could be effective tools to deal with the biodiversity crisis in the EU. However, in the last State of nature in the EU report (European Environment Agency, 2015), the conservation status of more than half of the species legally protected by the Directives were considered unfavourable, and studies on the effectiveness of N2000 came to very diverse conclusions (Davis et al., 2014; Popescu et al., 2014). Gruber et al. (2012), for instance, analysed the representation of N2000 Annex II species in the entire network according to their distribution ranges and found that most target species are well represented. However, they also found a strong bias in protection status with an overrepresentation of species with a large distribution range and an underrepresentation of species with a narrow distribution range. Additionally, a recent study by Hermoso et al. (2015) showed that N2000 fails to adequately cover freshwater biodiversity in Spain with some species never occurring within the network and less than 20% of the analysed species distribution ranges covered on average. Abellàn et al. (2011) found that the implementation of N2000 on the Iberian Peninsula will increase the protection status for raptor species. However, in terms of efficiency, defined as the proportion of nesting territories included within the system in relation to the surface of the system, the network always falls short of expectations. On the other hand, Pellissier et al. (2013) showed for France that the abundance of common, mostly non-target birds, i.e. species not listed in the Birds Directive, is higher within N2000 than outside. They concluded that the designation of N2000 sites for target species is also beneficial for non-target bird species. One of the few studies focusing on habitat types, conducted by Miklín and Čížek (2014), focused on open woodland habitats in Czech Republic, identifying habitat fragmentation and logging as the main causes for the disappearance of open woodland habitat types. Although open woodland habitat types are listed in the HD Annex I, even after the implementation of N2000 in the Czech Republic, these habitat types still disappear and the effectiveness of N2000 to protect them remains questionable. In conclusion, although the implementation of N2000 has positive effects on some taxonomic groups, the effectiveness of the network is still disputable.

Europe stands out as one of the areas in the world with the highest human-foot-print, indicating a high anthropogenic pressure on ecosystems (Venter et al., 2016). In densely populated areas, implementation of nature conservation areas is often biased towards remote or abandoned regions of low commercial interest (Margules & Pressey, 2000). Additionally, adding new areas to existing networks is often connected with high costs (Underwood et al., 2008) and is politically challenging. For N2000 this results in a static, inflexible network (Davis et al., 2014) including in some countries, such as for instance Germany, many small and unconnected protected areas (European Commission, 2016b). Especially for the 230 habitat types listed under Annex I, this could result in serious conservation problems due to for example habitat fragmentation, which is one of the key drivers for biodiversity loss (Haddad et al., 2015), and resulting edge-effects (Porensky & Young, 2013). An additional challenge for applying the HD is that there are no specific area targets set for each habitat type. This uncertainty may lead to misinterpretations regarding the degree of protection of individual habitat types and biased conservation efforts towards ‘charismatic’ habitats, as this has been shown for species conservation (Ducarme et al., 2013; Martín-López et al., 2009).

Despite these preconditions, including the fact that the conservation status of the listed habitat types is generally worse than that for species (Berry et al., 2016), surprisingly few studies have concentrated on the degree of protection of habitat types (but see Dimopoulos et al., 2006; Miklín & Čížek, 2014; Rosati et al., 2008) compared to the vast amount of studies focusing on single species or groups of species (Gruber et al., 2012; Hermoso et al., 2015; Hermoso & Kennard, 2012; Iojă et al., 2010; Kukkala et al., 2016; F. Lisón et al., 2013; Fulgencio Lisón et al., 2015; Pellissier et al., 2013; Rubio-Salcedo et al., 2013). A likely reason for this bias is the lack of information on N2000 habitat types outside the current N2000 network (Evans, 2006; Mücher et al., 2009). For example, the Mid-term review of the EU Biodiversity Strategy to 2020 (European Commission, 2015) identified the lack of information on forestry habitat types distribution outside N2000 as the reason for not significantly improving towards the Target 3b ‘Increase the contribution of forestry to maintaining and enhancing biodiversity’. As a consequence, studies addressing habitat protection under N2000 have either i) focussed only on a small number of habitats occuring in a country (Miklín & Čížek, 2014), or ii) assessed the status of habitat protection only in a small area of a country (Mikkonen & Moilanen, 2013).

Hence the two main objectives of our study were 1) to develop a workflow to map habitat types outside of N2000, information which is needed to evaluate the protection status of all habitat types considered in the HD at a country scale, and 2) to demonstrate how to identify areas to fill protection gaps effectively and cost-efficiently. To do so, we i) mapped the spatial distribution of habitat types at a 30 km^2^ resolution. Based on this information, ii) we performed a gap analysis to identify habitats that are currently not sufficiently protected and iii) used the conservation planning software Marxan to spatially optimize the current N2000 network testing for different targets, including a 17% target to account for Aichi target 11 (CBD 2017). We demonstrate this approach for Germany as a case study, because Germany is one of the most densely populated countries in the EU (Venter et al., 2016) which is especially challenging when establishing meaningful conservation areas. To the best of our knowledge, this is the first study that provides generic guidance on how to evaluate the representation of N2000 target habitats with a subsequent optimization to inform potential future management.

## 2. Methods

### 2.1 Study area

We selected Germany as the study area as covered by the German ordnance survey map based on TK25 quadrant map sheets (hereafter called TK25 map; Orchids, 2016). The TK25 map consists of 12028 grid cells and includes the terrestrial area of Germany, but excludes most of the marine territory, covering an area of approximately 360,000 km^2^. From the 230 habitats designated under N2000, 91 occur in Germany. The German N2000 network consists of more than 5300 mostly small (median area = 2 km^2^), single sites covering an area of about 80,000 km^2^.

### 2.2 Distribution of Natura 2000 habitats

To determine the gaps in habitat coverage under N2000, and to eventually identify priority areas for filling these gaps for those habitats that were not adequately covered, we needed maps with habitat-specific occurrence information. EU member states have to report N2000 habitat types within quadrants of 10 × 10 km (100 km^2^; www.bd.eionet.europa.eu). Because most of the German habitat mapping to date is based on the TK25 map, with 11 × 11 km grid cells (121 km^2^), Germany is currently using these quadrants to report N2000 habitat types. However, this information is too coarse for meaningful conservation planning, because i) these grid cells are too large to be considered for on-ground implementation by policy makers and stakeholders, and ii) fewer large areas mean less flexibility in the conservation planning process.

In order to refine the information given on grid cell level, we divided each grid cell into four quadrants of equal size (which is the unit we further used as planning unit (PU); 1 PU ~ 30 km^2^; total number of PUs = 48112; Fig. 1 C). Since the TK25 map only states whether or not a habitat occurs within a grid cell (and gives no information about its extent or location), we had to verify whether the habitat is located in one, two, three, or all four of the respective PUs. To do so, first we combined the divided TK25 map with a European Nature Information System (EUNIS) habitat class map (section 2.2.1 “Assignment of EUNIS habitat classes to N2000 habitats” and Fig. 1 D; European Environment Agency, 2016a). In a second step, we verified habitat occurrence on PU-level by using occurrence data of characteristic plant species (section 2.2.2 “Assignment of plant species to Natura 2000 habitats” and Fig. 1 E). With this process, we excluded PUs that have a high improbability of containing the target habitat. For further analyses, we used presence/absence data of habitats within a PU.

**Figure 1.**
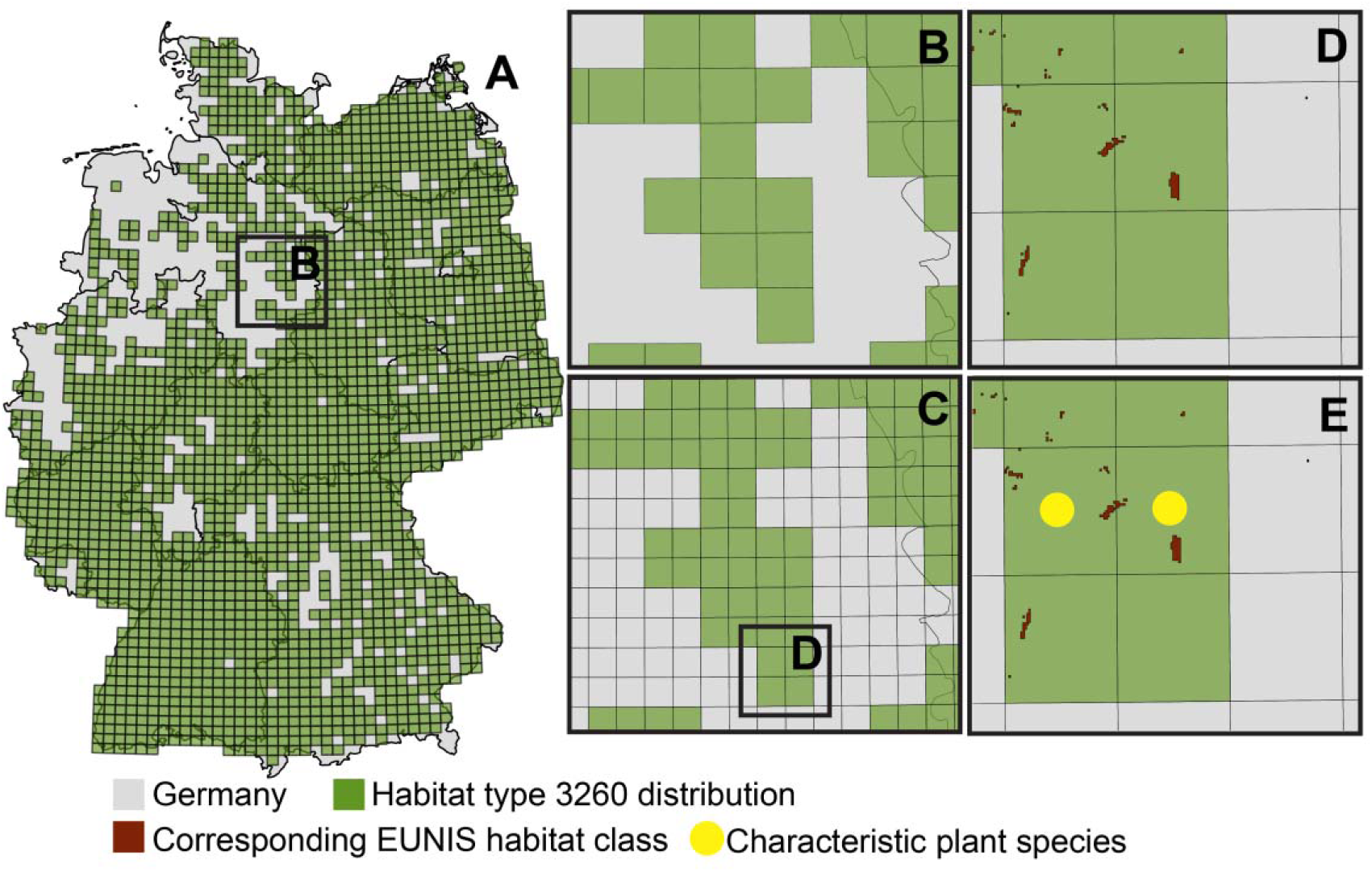
Example of PU-specification of N2000 habitats. A) Distribution of habitat 3260 “Water courses of plain to montane levels with the *Ranunculion fluitantis* and *Callitricho-Batrachion* vegetation” in Germany as reported by the German Federal Agency for Nature Conservation (BfN; BfN, 2013) on TK25 grid cell level (each grid cell is ~ 120 km^2^). B) Enlargement of a random area to better visualize the partitioning of TK25 grid cells into four equal quadrants (PUs; area ~ 30 km^2^) as seen in C. D) Single, partitioned TK25 grid cell combined with the EUNIS habitat class which accounts for habitat 3260. The lower right PU was excluded from further considerations of habitat 3260 occurrence, since it does not contain any EUNIS habitat class accounting for habitat 3260. E) Single, partitioned TK25 grid cell and the EUNIS habitat class combined with occurrence data of plant species characteristic for habitat 3260 (yellow circles). Since in the lower left PU not a single characteristic plant species accounting for habitat 3260 occurred, this PU was excluded from further considerations of potential habitat occurrence. In this example, habitat 3260 most likely occurs only in the upper two PUs.

#### 2.2.1 Assignment of EUNIS habitat classes to Natura 2000 habitats

The EUNIS habitat classification map offers habitat information with a spatial resolution of 100 × 100 m grid cells, which is much finer than the TK25 maps. However, its thematic resolution is coarser than the TK25 map considering only 40 different habitat classes in Germany (Supporting Information Table S1). Eight of these habitat classes belong to highly artificial areas and were excluded from further analyses (Table S1, asterisks). Within the remaining 32 habitat classes, only three different categories exist for forests, whereas in Annex I of N2000 18 different forest habitats are listed plus another two habitats, which are closely related to forests. To make use of the high spatial resolution of the EUNIS habitat classification map, we had to find matching combinations of the remaining 32 habitat classes to describe the 92 N2000 habitats occurring in Germany. Although recommendations on how habitat classes can be translated into N2000 habitats exist (European Environment Agency, 2018) we found this exercise challenging. To end up with a better coverage between the TK25 maps and the EUNIS habitat classes, first, we checked each EUNIS habitat class definition and searched for similarities in N2000 habitat definitions. Additionally, we used example pictures showing N2000 habitats from guidelines for field monitoring from different German Federal States and checked the reports from the German Federal Agency for Nature Conservation (BfN) for habitat-specific plant species. We used this information to find more similarities between plant species and EUNIS habitat classes. For example, N2000 habitat 3260 ‘Water courses of plain to montane levels with the *Ranunculion fluitantis* and *Callitricho-Batrachion* vegetation’ is described by the BfN as mainly consisting of streams and running waters, but also containing backwaters and near natural drains with natural vegetation (BfN, 2011). The recommended matching EUNIS land cover class is ‘Surface running waters’ (European Environment Agency, 2018). However, by using just this recommendation, we would have missed a high amount of PUs in which habitat 3260 occurs in Germany. Therefore, we additionally matched the EUNIS land cover class ‘Surface standing waters’ to the habitat 3260, since this class contains canals, man-made freshwater bodies, and reservoirs (European Environment Agency, 2016a). Moreover, because the EUNIS habitat classification is based on satellite images, and therefore does not indicate sub-surface habitats, we had to exclude N2000 habitat 8310, i.e. ‘Caves not open to the public’ from further analyses. We also excluded the habitats 1110, 1130, 1140, 1160, and 1170, as they belong to open sea and tidal areas. Additionally, these habitats mostly refer to areas which are not covered by the TK25 map.

#### 2.2.2 Assignment of plant species to Natura 2000 habitats

The BfN assigns a set of characteristic plant species based on plant communities to the majority of N2000 habitats occurring in Germany (i.e. to 83 of the 85 habitats that we included in this study). We used this information to further downscale the N2000 habitat distribution information provided by the BfN. We used occurrence data of plant species provided by www.floraweb.de, which reports information on the same grid cell level, i.e. PU level, as we used for our analyses (30 km^2^). The classification of these communities is based on syntaxonomy with the plant association as the highest syntaxonomic rank, which is further separated into alliances, orders, and classes (Pott, 1992). For each syntaxonomic rank, a so called character species or group of character species exists. Character species are plant species, which define a certain class, order, alliance, or association. Additionally, differential species are used to distinguish between classes, orders, alliances, or associations. That means that these plant species are used to separate two closely related classes, orders, alliances, or associations, which could have a similar set of character species. We checked habitat descriptions provided by the BfN as well as descriptions provided from the German Federal States for field habitat mapping to evaluate which plant community classes or orders are typical for each N2000 habitat. We used the lowest syntaxonomic rank possible. First, we included character species of the identified communities in our analyses. In case there were not enough character species, we used differential species to reach a number of ten plants per habitat (Table S2). In case of more than one plant community was recommended for a habitat, we selected an equal number of plant species across the communities for our analyses. Additionally, we checked for each habitat, if the species named as the character species by Pott (1995) also occured in the plant species list provided by the BfN. Only if that was true, we finally used a plant species for our downscaling approach.

### 2.3 Assessing habitat coverage

To assess the coverage of habitats under N2000, we first had to decide whether a PU can be considered as protected or not. To do so, we calculated the area of each PU covered by N2000 using the most current N2000 layer (European Environment Agency, 2016b). We built four scenarios using different thresholds of the percentage of a PU area covered by N2000 sites (90%, 75%, 50%, and 25%; scenario 90, 75, 50 and 25, respectively). Second, we tested each scenario for missing habitats, i.e. habitats that did not occur in any of the PUs considered as protected. Third, we evaluated whether a habitat can be considered as sufficiently covered. As a target, we used 17% of the total number of PUs where the respective habitat occurs, adopting Aichi target 11 for lack of specific area targets in the HD (for more information see Table S2). For example, if a habitat occured in 100 PUs, the habitat-specific target was set to 17 PUs. Fourth, to test whether the habitat-specific frequency of occurrence, measured as the amount of PUs in which a habitat occurs influences its coverage within N2000, we separated habitats in three groups: “rare” habitats found in less than 100 PUs, “intermediate” habitats in between 100 to 1000 PUs, and “common” habitats occuring in more than 1000 PUs.

### 2.4 Identification of priority areas using Marxan

To identify a minimum set of PUs to improve the representation of habitats in N2000, we used the conservation planning software Marxan (Ball et al., 2009). Marxan solves the so-called minimum-set problem by trying to represent all conservation features at the minimum cost, while attending to other spatial constrains like connectivity (Ball et al., 2009). In our study, we ran Marxan for five different scenarios. In scenarios 90, 75, 50, and 25, we locked-in the PUs, i.e. we forced Marxan to use the PUs in the planning exercise, which are covered by N2000 in more than 90%, 75%, 50%, or 25%, respectively (see above). In an additional free-choice scenario, we let Marxan choose from all PUs, apart from 173 PUs located at the North and Baltic Sea, which we locked-in (in all scenarios). In these PUs that cover mainly marine habitats and only partly terrestrial ones, N2000 sites were most likely implemented to protect habitats which we excluded from our analyses (see above). We used the average human footprint value (Sanderson et al., 2002; Venter et al., 2016), a measure of human impact in each PU (Fig. 2; Sedac, 2016) as a surrogate for cost (human footprint penalty = HFPP). Details of the Marxan analyses are elaborated in the SI.

**Figure 2.**
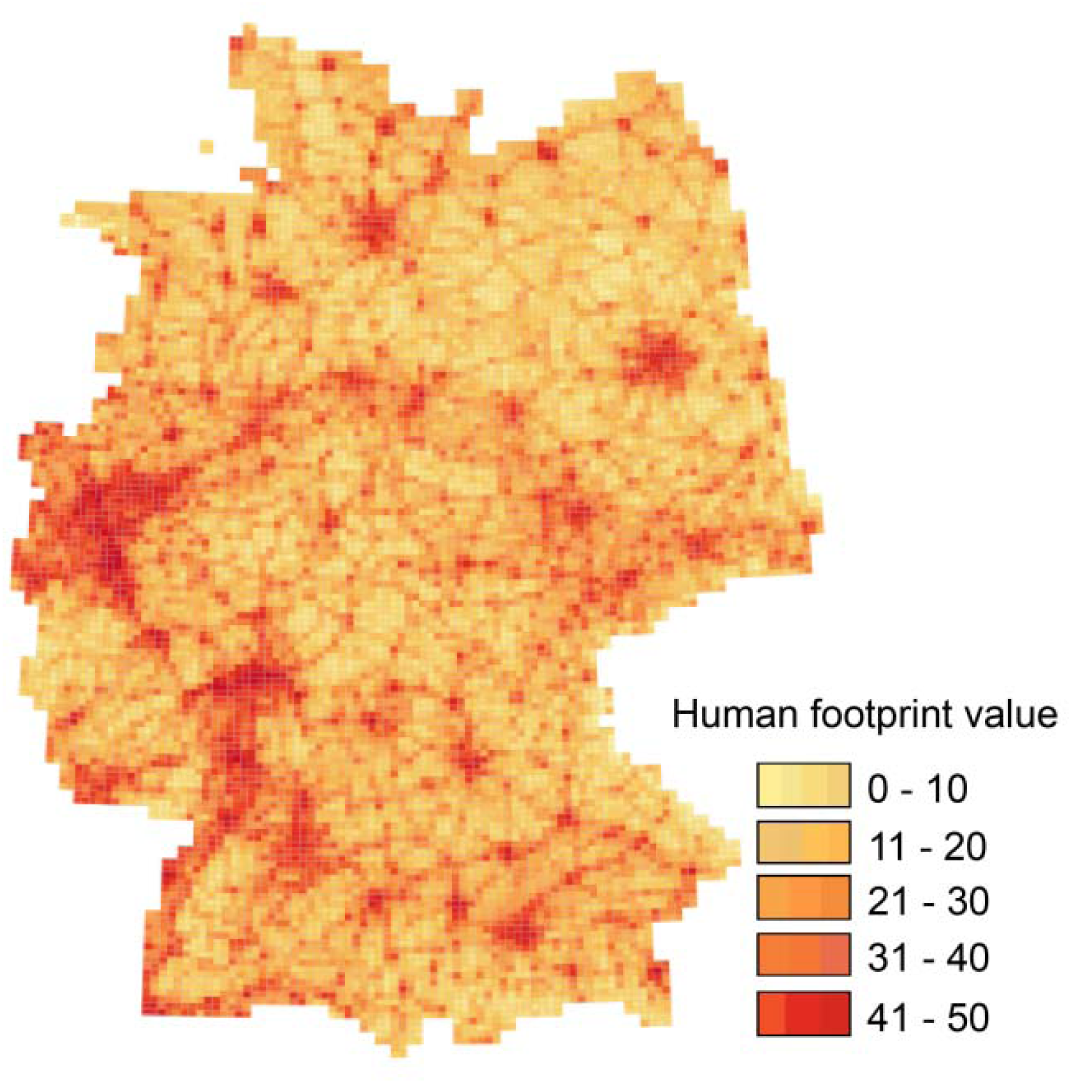
Human footprint map (HFP) of Germany. The HFP was used as a surrogate for highly humanized PUs for the Marxan analyses. A higher value indicates greater human impacts on a PU, i.e. a higher cost-penalty for adding a PU.

The result of high human footprint values in PUs is a reduced likelihood of these PUs to be chosen for the conservation network, since they are most likely heavily degraded and, therefore, of lower quality or very costly. As there are no specific area targets for habitats mentioned in the HD, we ran a sensitivity analyses for seven different area targets (1%, 5%, 10%, 15%, 17% (the Aichi Target 11), 20%, and 25%) calculated as percentages of the total number of PUs in which a habitat occurs, for all scenarios (see above). We set the species penalty factor (SPF) to 10 for all habitats, because we wanted to include all habitats with the same probability in our planning solution and calibrated the boundary length modifier (BLM) to two. For each target, we ran 100 replicates with 1,000,000 iterations each.

## 3. Results

### 3.1 Habitat coverage

Analysing habitat coverage revealed that the number of total missing habitats ranged from zero in scenario 25 to seven in scenario 90 (Table 1). Considering 17% of the total number of PUs in which a habitat occurs as a conservation target for each habitat, scenario 25 adequately covers 83 of 85 habitats, hence the two missing habitats were included in N2000 sites but not 306 covered enough to meat the 17% target. In scenario 50, the number of sufficiently protected habitat types declined to 37, and in scenario 90 it dropped to 15 habitats inadequately covered (Table 1).

**Table 1.**
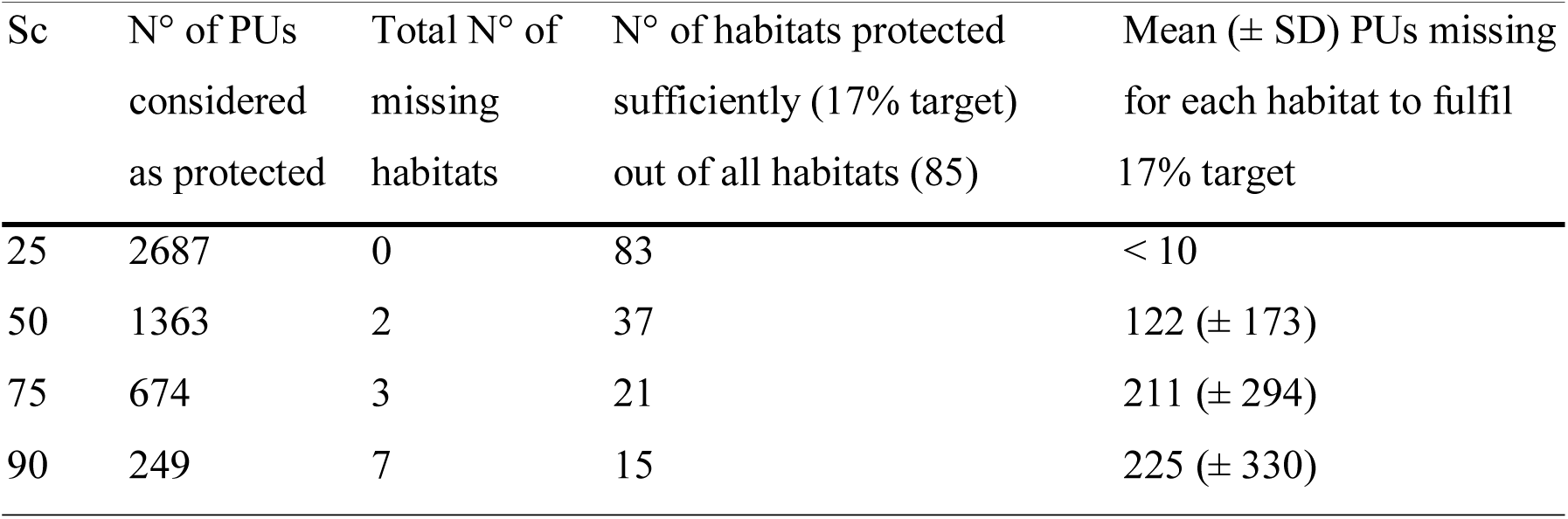
Overview of missing habitats in the four different scenarios. For each scenario, the number of PUs considered as protected, the number of total missing habitats, the number of habitats protected sufficiently when considering the 17% target (maximum 85 habitats), and the mean number of PUs which have to be additionally protected for each habitat to reach the 17% target. Sc = scenarios.

| Sc | N° of PUs considered as protected | Total N° of missing habitats | N° of habitats protected sufficiently (17% target) out of all habitats (85) | Mean (± SD) PUs missing for each habitat to fulfil 17% target |
| --- | --- | --- | --- | --- |
| 25 | 2687 | 0 | 83 | < 10 |
| 50 | 1363 | 2 | 37 | 122 (± 173) |
| 75 | 674 | 3 | 21 | 211 (± 294) |
| 90 | 249 | 7 | 15 | 225 (± 330) |

Although not statistically significant, there was a clear trend of decreasing coverage of N2000 habitats from rare to intermediate to common habitats for all scenarios (Fig. 3). In scenario 25, i.e. in the least restrictive scenario, all three habitat categories were covered by more than 17% of the total N2000 area with a mean proportion of 56.8 ± 21.3% (mean ± SD) for rare habitats, 44.8 ± 20.3% for intermediate, and 26.7 ± 4.7% for common habitats (Fig. 3). Contrarily in scenario 90, the proportional areas covered by N2000 were below 17% for all three categories (16.0 ± 17.8% for rare, 10.9 ± 12.3% for intermediate and 2.5 ± 2.0% for common habitats; Fig. 3). Scenarios 50 and 75 ranged in between these two scenarios, but showed the same trend of decreasing coverage with increasing number of PUs in which a habitat occurs (Fig. 3).

**Figure 3.**
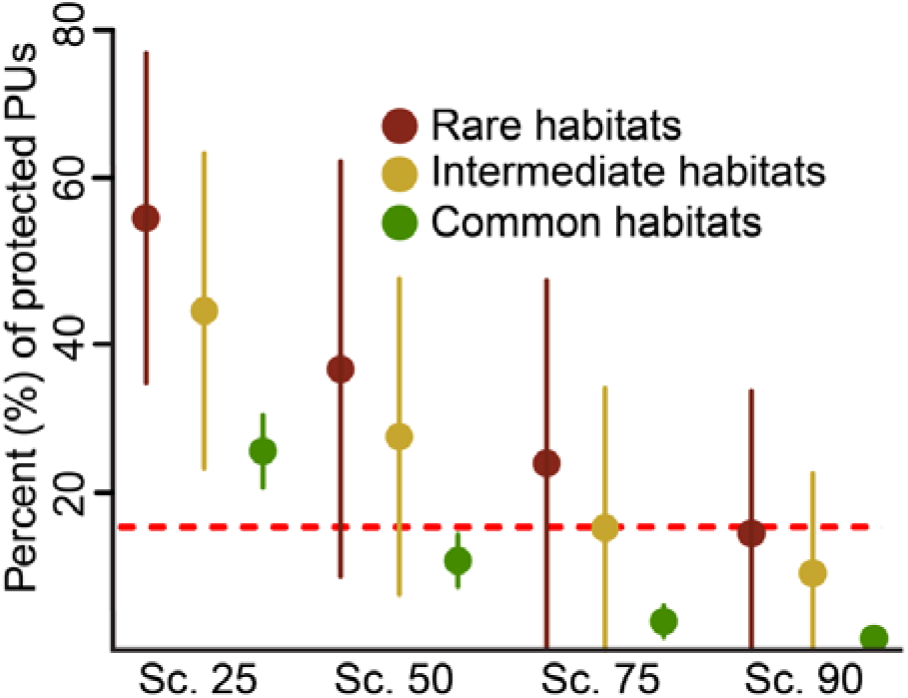
Covered protection of habitats under N2000. Representation of the three habitat categories (proportion covered by Natura 2000 sites, mean ± SD) in the current N2000 network based on the four different scenarios (25, 50, 75 and 90). Rare habitats occur in less than 100 PUs (n = 18), intermediate in a range of 100 to 1000 PUs (n = 39), and common habitats occur in more than 1000 PUs (n = 28). The red dashed line indicates the 17% target.

### 3.2 Identification of priority areas to fill habitat gaps using Marxan

To fill the identified habitat gaps, we ran a sensitivity analyses in Marxan for each of the four scenarios and the free choice scenario. The analyses revealed that the lower the threshold was set for a PU to be considered as protected, the more PUs were needed to reach a target and, therefore, the higher the total HFPP was (Fig. 4; Table 2; Table S2 with PUs per habitat in SI). This resulted in the highest total number of 2700 PUs needed in scenrio 25, and a total HFPP of 427.1, to fulfil the 17% target for all habitats. However, because N2000 in scenario 25 already consisted of 2687 PUs, only 13 PUs would have to be added (Table 2). Even for a 20% target, the number of additional PUs needed remains low (Fig. 4). Scenario 90 required the fewest PUs to be protected (1730) with the lowest HFPP (227.0; Table 2; Fig. 4). Numbers of PUs and HFPP in scenarios 50, 75, and free-choice ranged in between these two scenarios (Table 2). All scenarios considering the current N2000 network in the Marxan analyses, except scenario 90, were less efficient, i.e. needed more PUs to reach a certain target compared to the free-choice scenario, regardless the targets used (Table 2).

**Figure 4.**
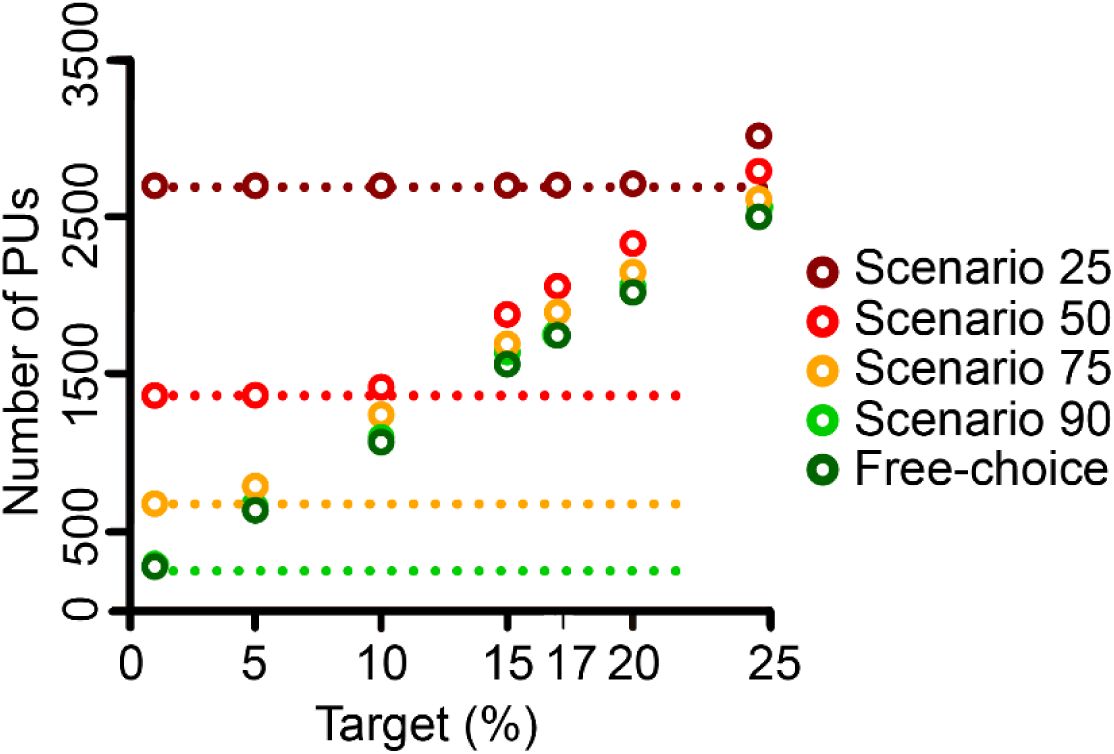
Sensitivity analysis for habitat target setting. Total number of PUs (i.e. locked-in PUs plus the ones identified by Marxan) needed to reach the 1%, 5%, 10%, 15%, 17%, 20%, or 25% target. Targets where calculated as a percentage of the maximum number of PUs in which a certain habitat occurs. For example, the 5%, 10%, 20% targets for a habitat which occurs in 100 PUs were 5, 10, and 20 PUs, respectively. Dashed lines indicate the number of PUs in each scenario, which were considered as initially protected, i.e. covered by N2000 and therefore locked-in for Marxan analyses (Table 1).

**Table 2.** Specifics on Marxan solutions. Detailed information about necessities to protect 17% of each N2000 habitat in each scenario. HFPP = human footprint penalty.

| Scenario | Locked-in PUs | N° of additional PUs needed to fulfil the 17% target | Total number of PUs | HFPP for additional PUs | Total HFPP |
| --- | --- | --- | --- | --- | --- |
| 25 | 2687 | 13 | 2700 | 1.32 | 427.1 |
| 50 | 1363 | 685 | 2058 | 91.05 | 284.0 |
| 75 | 674 | 1219 | 1893 | 155.89 | 243.4 |
| 90 | 249 | 1484 | 1733 | 193.43 | 227.0 |
| Free-choice | 173 | 1545 | 1745 | 203.75 | 233.93 |

However, in all five scenarios habitat targets could be reached; i.e. it was possible to represent the 85 habitats adequately by adding the identified PUs (Fig. 5.).

**Figure 5.**
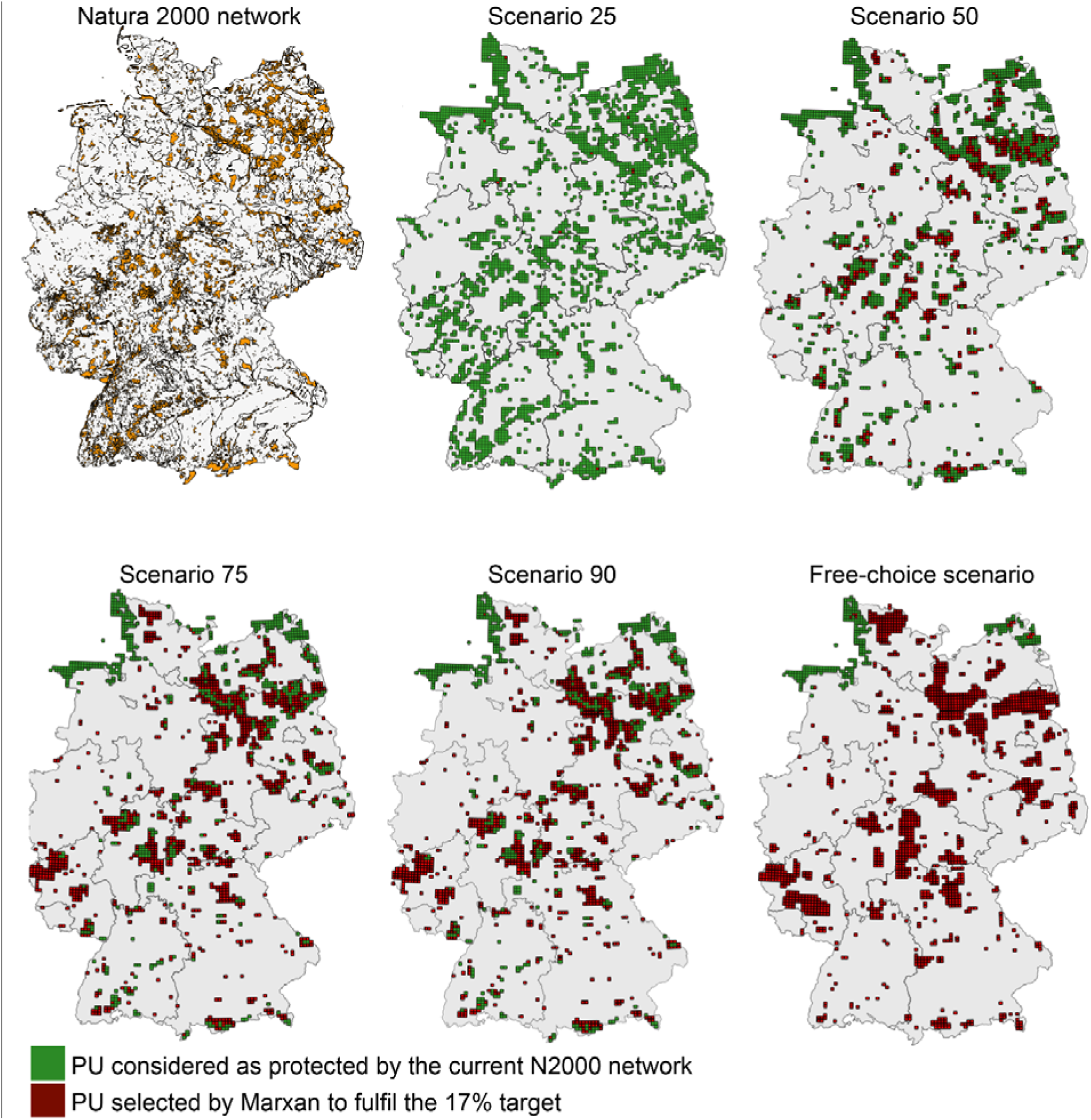
Best Marxan solutions for the five scenarios. Spatial distribution of PUs considered as protected (i.e. locked-in in the optimization analyses) and PUs selected by Marxan to fulfil the 17% target for all habitat types for the five scenarios. All scenarios met the 17% target for all habitat types (n = 85). Note that we locked-in PUs (green) being part of the coastal areas of North and Baltic Sea for the free-choice scenario (see Methods).

Finally, to evaluate if the current set of protected PUs under N2000 is the most effective way to cover habitats across all scenarios, we compared the spatial overlap of each scenario with the free-choice scenario (assuming that this scenario represents a highly efficient way of achieving the representation targets for all habitats; Fig. 5). We found an increase in spatial overlap when the threshold for considering a PU as protected increased. In scenario 25, for example, we found 22.2% of the initially protected PUs also being selected by Marxan in the free-choice scenario, whereas 35.5% of PUs overlapped in scenario 90. Scenarios 50 and 75 shared 26.4% and 29.1% of PUs with the free-choice scenario, respectively.

## 4. Discussion

Multiple conservation needs within each EU member state could easily lead to an ineffective distribution of nature conservation funds, i.e. protecting areas because of national interests, while these areas are not effective when considering country-specific targets (Hermoso et al., 2016; Hochkirch et al., 2013). Thinking of N2000 as a network covering 28 EU member states, different policies and opposing interests between them result in an additional challenge for establishing N2000 sites (cost)-efficiently (Kukkala et al., 2016). Hence, it is essential for N2000 and the related directives to assess their effectiveness and identify potential for future progress regularly. Our results indicate that, even if established in a highly populated country like Germany (Venter et al., 2016), N2000 could cope with multiple interests and potentially cover an unbiased representation of habitats. But, we also identified room for improvement for which the current N2000 network can build an excellent basis.

### 4.1 Missing habitats and underrepresentation of common habitats

Our gap analyses did not identify any missing habitats for scenario 25, and the number of missing habitats even remained low for the most conservative scenario (i.e. scenario 90 with seven missing habitats). This is a success for N2000, especially when considering that studies which focus on species protection in N2000 often found a varying number of species not being effectively protected (Gruber et al., 2012). Although N2000 is successful when considering the number of protected habitats, we found that common habitats are underrepresented compared to rare ones which are on average well covered (>17% of their distribution included in N2000). This is an interesting finding, since it is not in line with studies that consider species as conservation targets, which often found that rare and endemic species are underrepresented in conservation networks (de Novaes e Silva et al., 2014; Maiorano et al., 2015; Venter et al., 2014). The good representation of rare habitats under N2000 in Germany could be the result of the thresholds we used to consider for a PU to be protected. As we cannot evaluate for which habitats a N2000 site was initially designated, it is possible that a PU, which for example currently has 25% covered by N2000 to protect a large lake, concurrently protects one or several rare habitats that happen to occur in the same PU. The likelihood of erroneously considering a habitat adequately covered by N2000 increases the lower the threshold is for considering a PU as protected. Additionally, as N2000 sites cover on average 6% of a PU (with half of all PUs being covered between 0.1 and 21%), we labelled many PUs as not sufficiently protected and, therefore, likely missed a far amount of protected area. These disregarded PUs could theoretically increase the coverage of common habitats. However, as they are only covered by small patches of N2000, it is more likely that including these PUs would have even increased the representation of rare habitats. Finally, we found that the median area of a N2000 site in Germany is 2 km^2^ with 75% of the sites being smaller than 9 km^2^. This means that, despite Germany declaring by far the most N2000 sites (5206), the total area covered by these sites is comparably small (80,773 km^2^ total area including 55,170 km^2^ terrestrial area, compared to Spain covering the largest N2000 area with 1863 sites covering 222,142 km^2^ total area including 137,757 km^2^ terrestrial area, European Commission 2016b). Hence, the amount and/or the size of N2000 sites may be insufficient to protect common habitats adequately in Germany (Lawton et al., 2010), which is a common issue conservation networks are facing (Wilson et al., 2007).

### 4.2 Call for specific area targets to avoid over- or underrepresentation of habitats

It is impossible to evaluate progress in habitat protection in Europe without measurable targets (Juffe-Bignoli et al., 2016). The Aichi target 11, which aims to protect 17% of terrestrial area, could easily be applied for each habitat. In Germany (at least when considering scenario 25), only a small amount of areas would need to be added to the current N2000 network to reach a 17% target for all habitats. We used a percentage of the total distribution of habitats to set our targets to test the Aichi target 11 which treats all habitats equally, i.e. suggest to protect the same proportion of all of them. Additionally, we chose targets for the habitats in proportion to their representation within a member states’ territory as recommended by the HD (92/43/EEC, Article 3, paragraph 2). Using proportional targets, we may have emphasised common, large habitats (mainly meadows and forests in our study). Doing so, however, we accounted for the fact that large-area habitats, such as for example forests, simply need more space to be protected sufficiently (Evans, 2006; Lawton et al., 2010) which is crucial for endangered, large, terrestrial mammals (IUCN, 2016). Considering area targets for each habitat in the HD would ensure i) a balanced habitat protection within Europe, and ii) to evaluate the trajectory of habitat protection compared to a current baseline. Since the HFPP for reaching the 17% target in scenario 25 was low (Table 2), the necessary PUs are likely situated in natural and semi-natural areas. Hence, it could be expected that the costs for adding these PUs would also be low, which is important when planning for future conservation (Underwood et al., 2008).

### 4.3 N2000 compared to the free-choice scenario

The free-choice scenario always performed better in terms of minimizing the HFPP and the total number of PUs than most other scenarios, except from scenario 90 which performed similarly well. Additionally, we found that the spatial overlap with the free-choice scenario increased when the threshold for considering a PU as protected increased. The much higher number of PUs necessary to reach all the targets, together with the low spatial overlap of 22.2% when compared to the free-choice scenario, indicates that in scenario 25 a lot of PUs considered as protected do not contribute much to the conservation status of habitats. Although we found only 35.5 % spatial overlap between the free-choice scenario and scenario 90, the free-choice scenario did not perform better than scenario 90 in terms of total HFPP and number of PUs. This indicates that PUs which are currently covered 90% or more by N2000 sites, have the potential to build the basis for an efficient conservation network. However, since in scenario 90 only 250 PUs were initially considered as proteced i.e. locked-in, another 1500 PUs would be needed to fulfil the 17% targets for each habitat type.

### 4.4 Data robustness

As there is not yet a full monitoring done for N2000 habitat types outside of the N2000 network in Germany, the habitat information contained in the TK25 maps we used here is based on expert judgments and/or modeling approaches. Hence, our estimations of the distribution of N2000 habitat types have to be taken with a pinch of salt. Combining these data with the EUNIS habitat class map to downscale the resolution of the TK25 maps and to run a meaningful conservation exercise, adds another layer of uncertainty to our analyses. We reduced uncertainty as much as it is currently possible, by comparing the definitions of N2000 habitat types and EUNIS habitat classes using the official translation tables provided by EEA, by manually checking the spatial overlap between N2000 habitat types and EUNIS habitat classes, and by verifying our habitat distribution estimations with plant species distribution data.

### 4.5 Applying the workflow to other EU member states

To apply the proposed workflow to other EU member states, two main data sets are needed: 1) A country-specific EUNIS habitat class map, which is open source and available for all EU member states (http://eunis.eea.europa.eu/about). This data set provides information on the occurrence of habitat classes (see more information in the method’s section above) and recommendations on how those can be linked to N2000 habitat types. In case country-specific N2000 habitat maps exist (as the TK25 map for Germany), they can be combined with the EUNIS habitat class map (see above). 2) To verify and downscale the EUNIS habitat class map (or a combined EUNIS-N2000 habitat map), plant distribution data is needed. Similar data sets as the one we used for Germany can be found for other countries. Examples of such data sets are: The “Raster der floristischen Kartierung” (https://www.gbif.org/dataset/85ab1bf8-f762-11e1-a439-00145eb45e9a) providing plant distribution data with an identical resolution as the German one for Austria; the Species Observation System “Artportalen” with more than 53 million occurrence records from plants, animals, and funghi (https://www.artdatabanken.se/en/species-observations/) covering Sweden; or the information system “Anthos” (http://www.anthos.es/index.php?lang=en)with plant occurrence information for Portugal and Spain, containing more than 1.8 million occurrence points for plants across the Iberian Peninsula alone; to just name a few.

### 4.6 Future conservation management practices: quantitative vs. qualitative targets

Conservation targets, as e.g. the CBD Aichi target 11 that assess a purely quantitative progress in nature conservation efforts, have recently been challenged (Barnes, 2015; Barnes et al., 2018). Barnes (2018) argues that nature conservation efforts must focus on the quality of protected areas and not on their quantity. Our study shows that one consequence of using quantitative targets is identifying an overrepresentation of rare conservation features. This clearly contrasts the current focus on implementing protected areas for rare conservation features (Asaad et al., 2017). For habitats in the N2000 network, this focus finds its expression in the discrimination between habitats which are only listed for protection and those among them which are given priority, because they are particularly vulnerable. In our study, we found the highest proportion of priority habitat types in the rare habitat group (28%; 23% and 21% in the intermediate and common habitat groups, respectively). Hence, overrepresentation of rare N2000 habitat types is partly anchored in the HD itself.

Focusing on rare or threatened conservation features will always result in a ‘reaction-behaviour’ of conservation efforts; i.e. nature assets must be (or become) rare or threatened, before conservation resources will be allocated. This stands in contrast to recent evidence found for a dramatic decline in species groups which have not been in the focus of conservation efforts, because they were considered abundant and non-threatened (Hallmann et al., 2017; Inger et al., 2015). Hence, we conclude that for an effective biodiversity protection a mixture of both, qualitative and quantitative targets, is needed. Here “qualitative” expresses a general active management of local conservation actions, some of which are hard to quantify. However, such local actions are vital to make conservation effective in practice. Consequently, in the HD a discrimination between priority and non-priority habitats is helpful to protect rare and threatened habitats, which currently need most attention. Additionally, however, we argue for quantitative targets to foster conservation efforts towards common, large habitat types to protect them from becoming rare or threatened in the first place.

## 5. Conclusions

To cope with pressing biodiversity issues that Europe is currently facing, we need protected areas that allow to tackle the specificities, conservation efforts have to deal with nowadays: First, allowing for flexibility to cope with arising climate- and land-use changes (Struebig et al., 2015); second, accounting for cost-effectiveness when planning new or adjusting existing protected areas (Hermoso et al., 2016; Underwood et al., 2008); and third, striving for tangible targets, because effective nature conservation needs more than just a certain amount of protected area (Barnes, 2015; Maiorano et al., 2015). The workflow we presented here, based on puplicly available data sets, allows identifying gaps and biases in habitat representation within the N2000 network in Europe, with the subsequent prioritization of areas with high conservation value. In Germany, the N2000 network is effective in covering the HD Annex II habitats, but rare habitats are overrepresented compared to common habitats when using the same quantitative proportional targets for all of them. The German case study additionally suggests that especially highly protected PUs (PUs covered by more than 90% with N2000 sites) build an excellent basis towards an effective and efficient network that can fulfil its role as the world’s largest nature conservation network. Our results enhance the value of N2000 at preserving habitats in Germany, whereas the proposed generic workflow helps guiding future assessment and conservation efforts in Europe.

## Acknowledgements

This work was supported by the European Union’s Horizon 2020 research and innovation programme [grant number 642317 and the Marie Skłodowska-Curie grant number 748625], by the German Federal Ministry of Education and Research [grant number 01LN1320A], and by a Ramón y Cajal contract funded by the Spanish Ministry for Economy and Competitiveness [VH; grant number RYC-2013-13979]. We thank Dr. A. Bonn and two anonymous reviewers for helpful comments that significantly improved an earlier version of the manuscript and Irene Petrosillo for handling our manuscript.

## Supplementary data

Supplementary data related to this article can be found at…

**S1.** Description of how Marxan variables were set; Species penalty factor (SPF) and Boundary Length Modifier (BLM).

**Table S1. EUNIS habitat classes occurring in Germany with corresponding numerical codes.** Habitat classes marked with an asterisk were not used to describe Natura 2000 habitats (see explanation in text).

**Table S2. Natura 2000 habitat types and corresponding EUNIS habitat classes and plant species.** Additionally, the maximum number (n°) of PUs in which a habitat type occurs, the number of PUs covered by N2000 in each scenario (Sc. 90, Sc. 75, Sc. 50, and Sc. 25), and the number of PUs which have to be protected to reach the 17% Aichi target, are given. PU = planning unit.

## References

Abellán, M. D., Martinez, J. E., Palazon, J. A., Esteve, M. A., & Calvo, J. F. (2011). Efficiency of a protected-area network in a Mediterranean region: a multispecies assessment with raptors. Environmental Management, 47(5), 983–991. doi:10.1007/s00267-011-9640-5

Asaad, I., Lundquist, C. J., Erdmann, M. V., & Costello, M. J. (2017). Ecological criteria to identify areas for biodiversity conservation. Biological Conservation, 213, 309–316. doi:10.1016/j.biocon.2016.10.007

Ball, I. R., Possingham, H. P., & Watts, M. E. (2009). Marxan and Relatives: Software for Spatial Conservation Prioritization. Oxford: Oxford University Press.

Barnes, M. (2015). Aichi targets: Protect biodiversity, not just area. Nature, 526, 1.

Barnes, M., Glew, L., Wyborn, C., & Craigie, I. D. (2018). Preventing perverse outcomes from global protected area policy. Shifting the focus from quantity to quality to avoid perverse outcomes. PeerJ Preprints, No. e26486v1, 1–18. doi:10.7287/peerj.preprints.26486v1

Berry, P., Smith, A., Eales, R., Papadopoulou, L., Erhard, M., Meiner, A., et al. (2016). Mapping and assessing the condition of Europe’s ecosystems: progress and challenges-EEA contribution to the implementation of the EU Biodiversity Strategy to 2020.

BfN. (2011, accessed 12.01.2018). Fließgewässer der planaren bis montanen Stufe mit Vegetation des *Ranunculion fluitantis*. Retrieved from https://www.bfn.de/lrt/0316-typ3260.html

BfN. (2013, accessed 12.01.2018). Vollständige Berichtsdaten - Allgemeiner Berichtsteil (Anhang A). Retrieved from https://www.bfn.de/themen/natura-2000/berichte-monitoring/nationaler-ffh-bericht/berichtsdaten.html

Bunce, R. G. H., Bogers, M. M. B., Evans, D., Halada, L., Jongman, R. H. G., Mucher, C. A., et al. (2013). The significance of habitats as indicators of biodiversity and their links to species. Ecological Indicators, 33, 19–25. doi:10.1016/j.ecolind.2012.07.014

Butchart, S. H. M., Walpole, M., Collen, B., van Strien, A., Scharlemann, J. P. W., Almond, R. E. A., et al. (2010). Global Biodiversity: Indicators of recent declines. Science, 328, 1164–1168.

Cardinale, B. J., Duffy, J. E., Gonzalez, A., Hooper, D. U., Perrings, C., Venail, P., et al. (2012). Biodiversity loss and its impact on humanity. Nature, 486(7401), 59–67. doi:10.1038/nature11148

CBD. (2017, accessed 10.02.2017). Aichi Biodiversity Targets. Retrieved from https://www.cbd.int/sp/targets

Davis, M., Naumann, S., McFarland, K., Graf, A., & Evans, D. (2014). Literature review, the ecological effectiveness of the Natura 2000 Network. ETC/BD report to the EEA, 30 pp.

de Novaes e Silva, V., Pressey, R. L., Machado, R. B., VanDerWal, J., Wiederhecker, H. C., Werneck, F. P., et al. (2014). Formulating conservation targets for a gap analysis of endemic lizards in a biodiversity hotspot. Biological Conservation, 180, 1–10. doi:10.1016/j.biocon.2014.09.016

Diaz, S., Fargione, J., Chapin, F. S., 3rd, & Tilman, D. (2006). Biodiversity loss threatens human well-being. PLoS Biology, 4(8), 1300–1305. doi:10.1371/journal.pbio.0040277

Dimopoulos, P., Bergmeier, E., & Fischer, P. (2006). Natura 2000 habitat types of Greece evaluated in the light of distribution, threat and responsibility. Biology & Environment: Proceedings of the Royal Irish Academy, 106(3), 175–187. doi:10.3318/bioe.2006.106.3.175

Ducarme, F., Luque, G. M., & Courchamp, F. (2013). What are “charismatic species” for conservation biologists? BioSciences Master Reviews, 10(2013), 1–8.

European Commission. (2015). Report from the commission to the European Parlament and the council - The mid-term review of the EU Biodiversity Strategy to 2020. http://www.ipex.eu/IPEXL-WEB/dossier/document/COM20150478.do

European Commission. (2016a, accessed 25.10.2016). Natura 2000- In a nutshell. Retrieved from http://ec.europa.eu/environment/nature/natura2000/index_en.htm

European Commission. (2016b, accessed 25.10.2016). Natura 2000 Barometer. Retrieved from http://ec.europa.eu/environment/nature/natura2000/barometer/index_en.html

European Environment Agency. (2015). State of nature in the EU. Results from reporting under the nature directives 2007-2012. https://www.eea.europa.eu/publications/state-of-nature-in-the-eu/at_download/file

European Environment Agency. (2016a, accessed 25.08.2016). EUNIS habitat types. Retrieved from https://www.eea.europa.eu/data-and-maps/data/ecosystem-types-of-europe/

European Environment Agency. (2016b, accessed 25.08.2016). Natura 2000 data- the European network of protected sites. Retrieved from http://www.eea.europa.eu/data-and-maps/data/natura-7#tab-gis-data

European Environment Agency. (2018, accessed 12.01.2018). Crosswalk EUNIS 2007 and Annex I 2008. Retrieved from https://www.eea.europa.eu/data-and-maps/data/eunis-habitat-classification#tab-documents

Evans, D. (2006). The habitats of the European Union Habitats Directive. Biology & Environment: Proceedings of the Royal Irish Academy, 106(3), 167–173. doi:10.3318/bioe.2006.106.3.167

Gruber, B., Evans, D., Henle, K., Bauch, B., Schmeller, D., Dziock, F., et al. (2012). “Mind the gap!” — How well does Natura 2000 cover species of European interest? Nature Conservation, 3, 45–62. doi:10.3897/natureconservation.3.3732

Haddad, N. M., Brudvig, L. A., Clobert, J., Davies, K. F., Gonzalez, A., Holt, R. D., et al. (2015). Habitat fragmentation and its lasting impact on Earth’s ecosystems. Science Advances, 1(e1500052), 1–10.

Hallmann, C. A., Sorg, M., Jongejans, E., Siepel, H., Hofland, N., Schwan, H., et al. (2017). More than 75 percent decline over 27 years in total flying insect biomass in protected areas. PloS ONE, 12(10), e0185809. doi:10.1371/journal.pone.0185809

Hermoso, V., Clavero, M., Villero, D., & Brotons, L. (2016). EU’s conservation efforts need more strategic investment to meet continental conservation needs. Conservation Letters, 10(2), 231–237. doi:10.1111/conl.12248

Hermoso, V., Filipe, A. F., Seguardo, P., & Beja, P. (2015). Effectiveness of a large reserve network in protecting freshwater biodiversity: a test for the Iberian Peninsula. Freshwater Biology, 60, 698–710. doi:10.1111/fwb.12519

Hermoso, V., & Kennard, M. J. (2012). Uncertainty in coarse conservation assessments hinders the efficient achievement of conservation goals. Biological Conservation, 147(1), 52–59. doi:10.1016/j.biocon.2012.01.020

Hochkirch, A., Schmitt, T., Beninde, J., Hiery, M., Kinitz, T., Kirschey, J., et al. (2013). Europe needs a new vision for a Natura 2020 network. Conservation Letters, 6(6), 462–467. doi: 10.1111/conl.12006

Hodge, I., Hauck, J., & Bonn, A. (2015). The alignment of agricultural and nature conservation policies in the European Union. Conservation Biology, 29(4), 996–1005. doi: 10.1111/cobi.12531

Hortal, J., Triantis, K. A., Meiri, S., Thebault, E., & Sfenthourakis, S. (2009). Island species richness increases with habitat diversity. The American Naturalist, 174(6), E205–217. doi:10.1086/645085

Inger, R., Gregory, R., Duffy, J. P., Stott, I., Vorisek, P., & Gaston, K. J. (2015). Common European birds are declining rapidly while less abundant species’ numbers are rising. Ecology Letters, 18(1), 28–36. doi:10.1111/ele. 12387

Iojă, C. I., Pătroescu, M., Rozylowicz, L., Popescu, V. D., Vergheleţ, M., Zotta, M. I., et al. (2010). The efficacy of Romania’s protected areas network in conserving biodiversity. Biological Conservation, 143(11), 2468–2476. doi:10.1016/j.biocon.2010.06.013

IUCN. (2016, accessed 10.02.2017). The IUCN Red List of Threatened Species. Retrieved from http://www.iucnredlist.org/initiatives/mammals/analysis/habitat

IUCN. (2017, accessed 10.02.2017). New Red List reveals European habitats under threat. Retrieved from https://www.iucn.org/news/new-red-list-reveals-european-habitats-under-threat

Juffe-Bignoli, D., Harrison, I., Butchart, S. H. M., Flitcroft, R., Hermoso, V., Jonas, H., et al. (2016). Achieving Aichi Biodiversity Target 11 to improve the performance of protected areas and conserve freshwater biodiversity. Aquatic Conservation: Marine and Freshwater Ecosystems, 26, 133–151. doi:10.1002/aqc.2638

Kukkala, A. S., Arponen, A., Maiorano, L., Moilanen, A., Thuiller, W., Toivonen, T., et al. (2016). Matches and mismatches between national and EU-wide priorities: Examining the Natura 2000 network in vertebrate species conservation. Biological Conservation, 198, 193–201. doi:10.1016/j.biocon.2016.04.016

Lawton, J. H., Brotherton, P. N. M., Brown, V. K., Elphick, C., Fitter, A. H., Forshaw, J., et al. (2010). Making Space for Nature: A review of England’s Wildlife Sites and Ecological Network.

Lisón, F., Palazón, J. A., & Calvo, J. F. (2013). Effectiveness of the Natura 2000 Network for the conservation of cave-dwelling bats in a Mediterranean region. Animal Conservation, 16(5), 528–537. doi:10.1111/acv.12025

Lisón, F., Sánchez-Fernández, D., & Calvo, J. F. (2015). Are species listed in the Annex II of the Habitats Directive better represented in Natura 2000 network than the remaining species? A test using Spanish bats. Biodiversity and Conservation, 24(10), 2459–2473. doi:10.1007/s10531-015-0937-1

Maiorano, L., Amori, G., Montemaggiori, A., Rondinini, C., Santini, L., Saura, S., et al. (2015). On how much biodiversity is covered in Europe by national protected areas and by the Natura 2000 network: insights from terrestrial vertebrates. Conservation Biology, 29(4), 986–995. doi:10.1111/cobi.12535

Margules, C. R., & Pressey, R. L. (2000). Systematic conservation planning. Nature, 405, 143–253.

Martín-López, B., Montes, C., Ramírez, L., & Benayas, J. (2009). What drives policy decision-making related to species conservation? Biological Conservation, 142(7), 1370–1380. doi:10.1016/j.biocon.2009.01.030

Mikkonen, N., & Moilanen, A. (2013). Identification of top priority areas and management landscapes from a national Natura 2000 network. Environmental Science & Policy, 27, 11–20. doi:10.1016/j.envsci.2012.10.022

Miklín, J., & Čížek, L. (2014). Erasing a European biodiversity hot-spot: Open woodlands, veteran trees and mature forests succumb to forestry intensification, succession, and logging in a UNESCO Biosphere Reserve. Journal for Nature Conservation, 22(1), 35–41. doi:10.1016/j.jnc.2013.08.002

Mücher, C. A., Hennekens, S. M., Bunce, R. G. H., Schaminée, J. H. J., & Schaepman, M. E. (2009). Modelling the spatial distribution of Natura 2000 habitats across Europe. Landscape and Urban Planning, 92(2), 148–159. doi:10.1016/j.landurbplan.2009.04.003

Orchids. (2016, accessed 25.08.2016). KoordinatenErmittler. Retrieved from http://www.orchids.de/haynold/tkq/KoordinatenErmittler.php

Pellissier, V., Touroult, J., Julliard, R., Siblet, J. P., & Jiguet, F. (2013). Assessing the Natura 2000 network with a common breeding birds survey. Animal Conservation, 16(5), 566–574. doi:10.1111/acv.12030

Popescu, V. D., Rozylowicz, L., Niculae, I. M., Cucu, A. L., & Hartel, T. (2014). Species, habitats, society: an evaluation of research supporting EU’s Natura 2000 network. PloS ONE, 9(11), e113648. doi:10.1371/journal.pone.0113648

Porensky, L. M., & Young, T. P. (2013). Edge-effect interactions in fragmented and patchy landscapes. Conservation Biology, 27(3), 509–519. doi:10.1111/cobi.12042

Pott, R. (1992). Die Pflanzengesellschaften Deutschlands—Verlag Eugen Ulmer: Stuttgart.

Rosati, L., Marignani, M., & Blasi, C. (2008). A gap analysis comparing Natura 2000 vs National Protected Area network with potential natural vegetation. Community Ecology, 9(2), 147–154. doi:10.1556/ComEc.9.2008.2.3

Rubio-Salcedo, M., Martínez, I., Carreño, F., & Escudero, A. (2013). Poor effectiveness of the Natura 2000 network protecting Mediterranean lichen species. Journal for Nature Conservation, 21(1), 1–9. doi:10.1016/j.jnc.2012.06.001

Sanderson, E. W., Jaiteh, M., Levy, M. A., Redford, K. H., Wannebo, A. V., & Woolmer, G. (2002). The human footprint and the last of the wild. BioScience, 52(10), 891–904.

Sedac. (2016). Human footprint. Retrieved from http://sedac.ciesin.columbia.edu/

Struebig, M. J., Wilting, A., Gaveau, D. L., Meijaard, E., Smith, R. J., Borneo Mammal Distribution, C., et al. (2015). Targeted conservation to safeguard a biodiversity hotspot from climate and land-cover change. Current Biology, 25(3), 372–378. doi:10.1016/j.cub.2014.11.067

Underwood, E. C., Shaw, M. R., Wilson, K. A., Kareiva, P., Klausmeyer, K. R., McBride, M. F., et al. (2008). Protecting biodiversity when money matters: maximizing return on investment. PloS ONE, 3(1), e1515. doi:10.1371/journal.pone.0001515

Venter, O., Fuller, R. A., Segan, D. B., Carwardine, J., Brooks, T., Butchart, S. H., et al. (2014). Targeting global protected area expansion for imperiled biodiversity. PLoS Biology, 12(6), e1001891. doi:10.1371/journal.pbio.1001891

Venter, O., Sanderson, E. W., Magrach, A., Allan, J. R., Beher, J., Jones, K. R., et al. (2016). Sixteen years of change in the global terrestrial human footprint and implications for biodiversity conservation. Nature Communications, 7, 12558. doi:10.1038/ncomms12558

Wilson, K. A., Underwood, E. C., Morrison, S. A., Klausmeyer, K. R., Murdoch, W. W., Reyers, B., et al. (2007). Conserving biodiversity efficiently: what to do, where, and when. PLoSBiology, 5(9), e223. doi:10.1371/journal.pbio.00502233.

